# Reward-associated configural cues elicit theta oscillations of rat retrosplenial neurons phase-locked to LFP theta cycles

**DOI:** 10.1101/2020.09.13.295444

**Authors:** Masashi Yoshida, Choijiljav Chinzorig, Jumpei Matsumoto, Hiroshi Nishimaru, Mitsuaki Yamazaki, Taketoshi Ono, Hisao Nishijo

**Author notes:** These authors contributed equally to this paper (Masashi Yoshida, Choijiljav Chinzorig). Correspondence to Dr. Hisao Nishijo, System Emotional Science, Faculty of Medicine, University of Toyama, Toyama 930-0194, Japan.

## Abstract

Previous behavioral studies implicated the retrosplenial cortex (RSC) in stimulus-stimulus associations, and also in retrieval of remote associative memory based on EEG theta oscillations. To investigate the neural mechanisms underlying these processes, RSC neurons and local field potentials (LFPs) were simultaneously recorded from well-trained rats performing a cue-reward association task. In the task, simultaneous presentation of two multimodal conditioned stimuli (configural CSs) predicted a reward outcome opposite to that associated with individual presentation of each elemental CS. Here, we show neurophysiological evidence that the RSC is involved in stimulus-stimulus association where configural CSs are discriminated from each elementary CS that is a constituent of the configural CSs, and that memory retrieval of rewarding CSs is associated with theta oscillation of RSC neurons during CS presentation, which is phase-locked to LFP theta cycles.

## Introduction

The retrosplenial cortex (RSC) is a functional hub where unimodal, multimodal and cognitive information converges: the RSC has reciprocal connections with the visual, parietal, motor, parahippocampal and prefrontal cortices, and hippocampal formation, and receives afferents from the auditory cortex [Vogt and Miller, 1983; van Groen and Wyss, 1990, 1992; Kerr et al., 2007; Kobayashi and Amaral, 2003; Vann et al., 2009; Mitchell et al., 2018; Todd et al., 2016, 2019]. Consistent with these diverse inputs, RSC neurons are reported to be have spatially correlated activity (Chen et al. 1994a,b; Cho and Sharp 2001; Jacob et al. 2017; Vann et al. 2009; Alexander and Nitz, 2017; Milczarek et al., 2018; Chinzorig et al., 2020; Mao et al., 2020). But lesion studies have also reported RSC involvement in processing of nonspatial information. RSC lesions impaired acquisition of serial or compound feature negative discrimination tasks (Keene and Bucci, 2008; Robinson et al., 2011) and a negative patterning discrimination task (Sutherland and Hoesing, 1993), in which serial or simultaneous presentation of a tone and a light was associated with nonreward, whereas the tone alone and/or the light alone predicted reward. Furthermore, chemogenetic silencing of RSC neurons induced acquisition deficits of tone-light association in sensory preconditioning, with serial presentation of tone and light cues (Robinson et al., 2014). These behavioral studies suggest that the RSC is involved in stimulus-stimulus associations (Robinson et al., 2011), wherein multiple stimuli were perceived together as a configural stimulus, as opposed to coincident individual elementary stimuli (Rudy and Sutherland, 1995). However, neurophysiological evidence indicating differential neuronal responses to elementary and configural CSs in the RSC is lacking.

Recent behavioral studies reported that the RSC is also implicated in storing and expression or retrieval of remote memory of nonspatial cues associated with reinforcement (Todd et al., 2016; Jiang et al., 2018; Fournier et al., 2019a). Theta oscillation is implicated in neurophysiological mechanisms for retrieval of remote memory: it reflects memory search and information transfer between the hippocampal formation and the cortical areas such as the RSC (Hasselmo, 2002; Fell and Axmacher, 2011; Burke et al., 2014). A human EEG study reported that timing of memory retrieval was synchronized with hippocampal theta cycles (Kerrén et al., 2018), consistent with a computational model where encoding and retrieval occurs on opposite phases of theta (Hasselmo et el., 2002). Furthermore, EEG theta oscillations occurred in the RSC 500-50 ms before retrieval of items (Kerrén et al., 2018). These findings suggest that RSC neuronal activity for long term memory is phase-locked to specific phases of theta cycles of local field potentials (LFPs).

In the present study, rats were well trained in a cue-reward association task to form long term association memory (Fig. 1). In the task, configural CSs consisting of two simultaneously presented elementary CSs predicted reward outcome opposite to that associated with each elementary CS. Thus, the rats were required to discriminate among six CSs associated with reward (sucrose solution) or nonreward (Table 1), and could obtain reward if they licked a tube after the rewarding CSs. Based on the previous behavioral studies, we hypothesized that RSC neurons would discriminate configural CSs from each elementary CS that is a constituent of the configural CSs. Second, we also hypothesized that if RSC neurons are involved in long term association memory, they would show theta oscillation phase-locked to LFP theta cycles in response to well-trained familiar CSs. Here, we show that RSC neurons code cue-reward association, and that response characteristics of these RSC neurons are consistent with the above two hypotheses.

**Table 1.** A list of conditioned stimuli (CS) associated with reinforcement (sucrose) or no reinforcement.

| Conditioned stimuli | Reinforcement |
| --- | --- |
| Elemental stimuli |  |
| Auditory stimuli |  |
| Tone 1 (2860 Hz) | Reinforcement (Sucrose) |
| Tone 2 (530 Hz) | No reinforcement |
| Visual stimuli |  |
| Light 1 (right) | Reinforcement (Sucrose) |
| Light 2 (left) | No reinforcement |
| Configural stimuli |  |
| Tone 1 + Light 1 | No reinforcement |
| Tone 2 + Light 2 | Reinforcement (Sucrose) |

**Figure 1.**
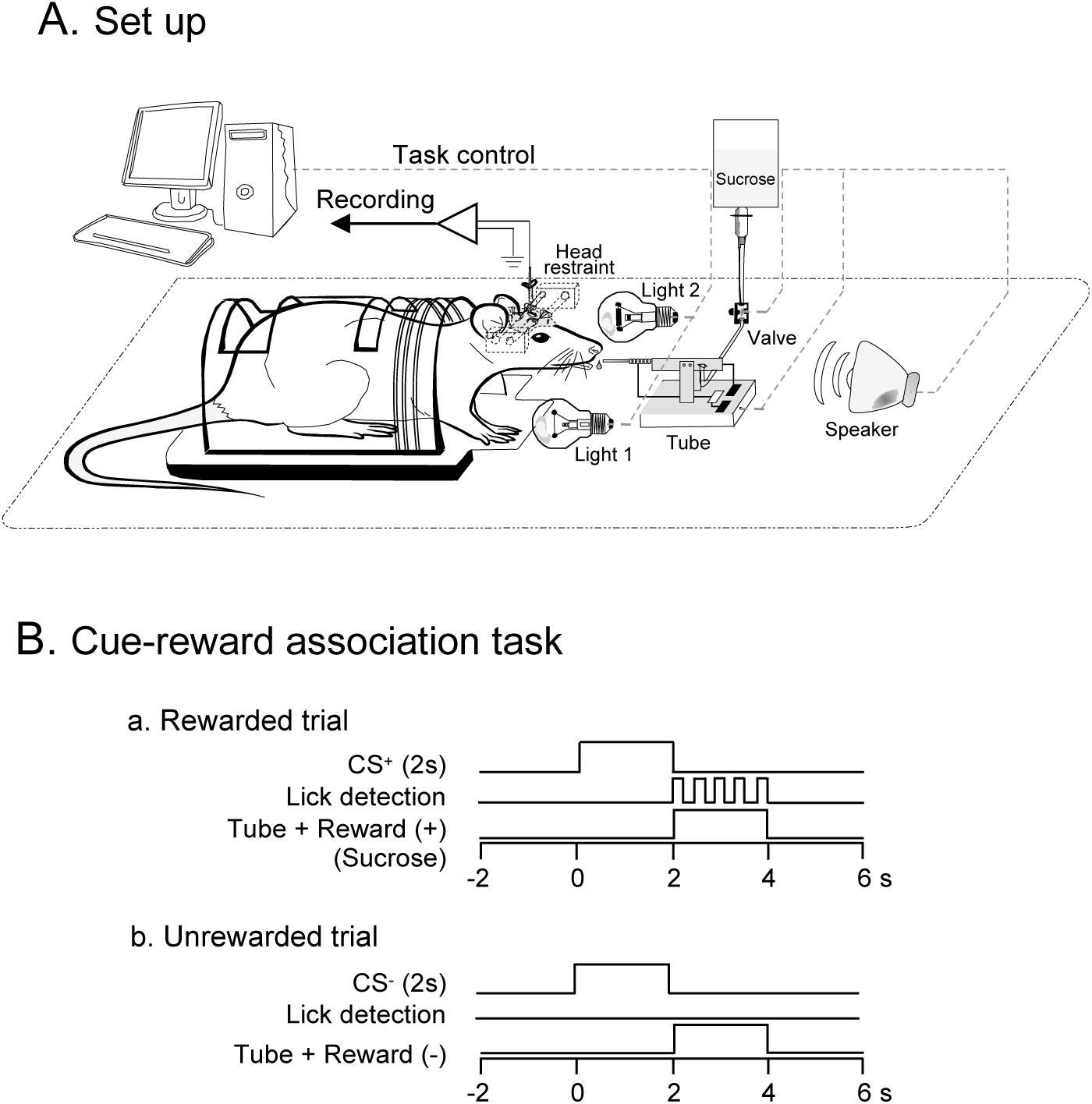
Schematic diagram of the experimental system. A, Schema of the experimental setup. Rats were painlessly placed in a stereotaxic apparatus by artificial ear bars (not shown). A speaker and two lights were placed in front of the rat to deliver conditioned stimuli (CSs). A movable tube was placed near its mouth. B, Time course of the cue-reward association task. (a) In the rewarded trials, one of CSs (tone, light, or configural stimuli) associated with reward was presented for 2 s before a tube was placed close to the rat’s mouth. Licking of the tube was detected by a touch sensor triggered by the tongue. (b) In the unrewarded trials, CS and tube protrusion were similarly presented to the rat. However, the reward was not delivered, and, as shown in this example, the rat did not usually lick the tube.

## Results

### RSC neuron responses in the task

Of 234 RSC neurons, 188 (80.3% 188/234) responded to at least one CS (CS-responsive neurons). A total of 215 neurons (91.9%, 215/234) showed excitatory responses during at least one of the six tube protrusion periods following the six CSs. Of the 215 tube period-responsive neurons, 33 responded only during the tube period, and 182 responded during both tube and CS periods. In the present study, we focused RSC neuronal activity in response to CSs, and Table 2 summarizes the response patterns of these 188 CS-responsive neurons). Of 188 CS-Responsive neurons 70 (29.9% 70/234) responded nondifferentially to the CSs (CS-nondifferential neurons), while 118 (50.4%, 118/234) responded differentially to the CSs. Of the 118 CS-differential neurons, 53 (22.5%, 53/234) responded stronger to all of the rewarding CSs than all of the unrewarding CSs (CS^+^-selective neurons).

**Table 2.**
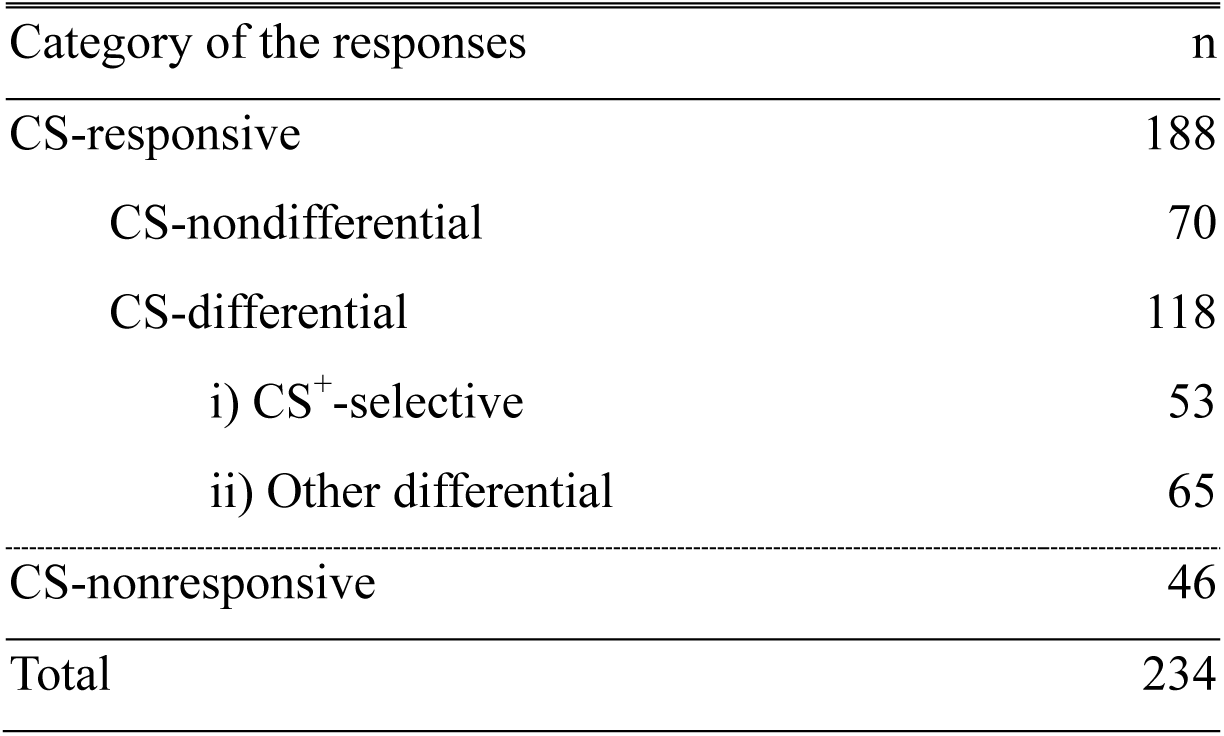
Category of the RSC neuronal response types.

| Category of the responses | n |
| --- | --- |
| CS-responsive | 188 |
| CS-nondifferential | 70 |
| CS-differential | 118 |
| i) CS<sup>+</sup>-selective | 53 |
| ii) Other differential | 65 |
| CS-nonresponsive | 46 |
| Total | 234 |

Figures 2A shows examples of superimposed waveforms of a representative RSC neuron and its autocorrelogram. The autocorrelogram indicated that the refractory periods were greater than 2.0 ms, suggesting that single unit activity was recorded. Figure 3 shows an example of a CS^+^-selective response (same neuron shown in Fig. 2). The RSC neuron showed excitatory responses to Tone1 and Light1 associated with reward (A, B), whereas the same RSC neuron did not respond to the configural stimulus consisting of Tone1 and Light1 associated with nonreward (C). Furthermore, although the RSC neuron did not respond to Tone2 and Light2 associated with nonreward (D, E), the neuron showed excitatory responses to the configural stimulus consisting of Tone2 and Light2 associated with reward (F). Figure 2B compares the response magnitudes to the six CSs for this RSC neuron. A one-way ANOVA showed a significant main effect of cue type [F (5, 24) = 32.386, p= 0.0001]. The neuron responded more strongly to all of the rewarding CSs (Tone1, Light1 and Tone2+Light2) than all of the unrewarding CSs (Tone2, Light2, and Tone1+Light1) (Bonferroni post-hoc test, p<0.01). Figure 2C shows mean response magnitudes of the all CS^+^-selective neurons (n=53) to the CSs. A statistical analysis indicated a significant main effect of cue type [F(5,312) = 81.570, p=0.0001]. These neurons responded more strongly to the rewarding CSs than the unrewarding CSs (Bonferroni post-hoc test, p<0.0001).

**Figure 2.**
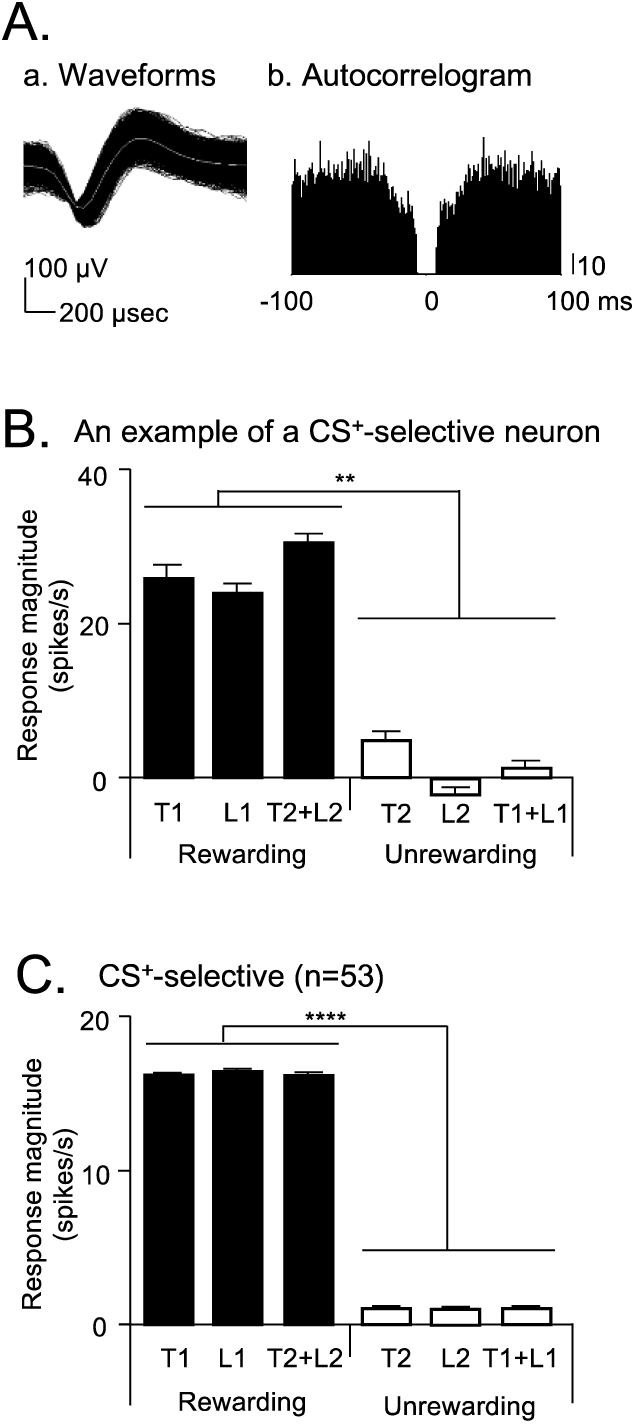
An example of a CS^+^-selective neuron (A, B) and mean CS^+^-selective responses (C). A, Identification of an RSC neuron. (a) Superimposed waveforms. (b) Autocorrelogram of this neuron. Bin width = 1 ms, where bin counts were divided by the number of spikes in the spike train. B, Histogram of response magnitudes during presentation of each CS. C, Responses in the 53 CS^+^-selective neurons. T1, Tone1; L1, Light1; T1+L1, Tone1+Light1; T2, Tone2; L2, Light2; T2+L2, Tone2+Light2. **, p < 0.01 (Bonferroni test). ***, p < 0.0001 (Bonferroni test). Histograms indicate mean ± standard error of the mean (SEM).

**Figure 3.**
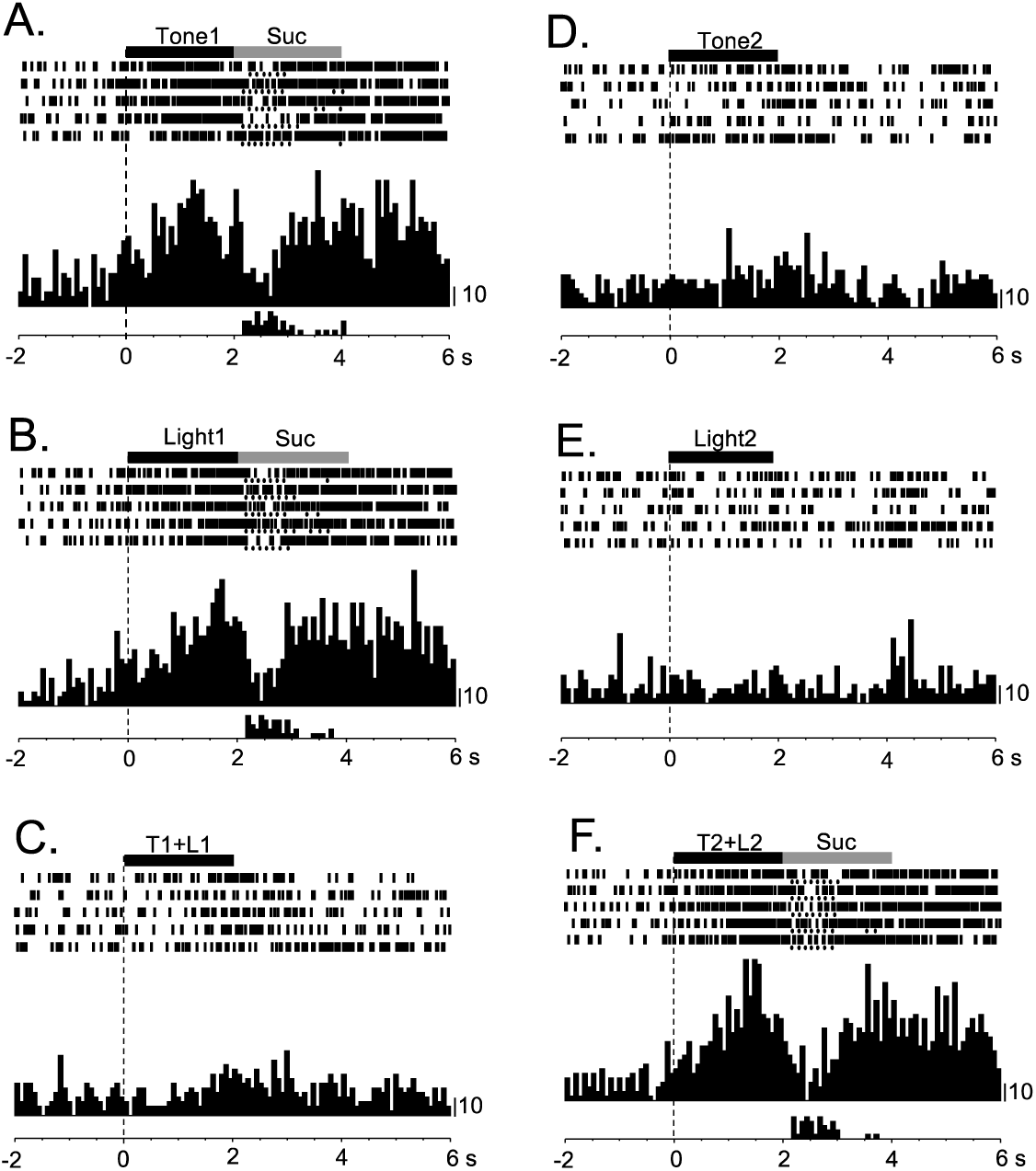
A CS^+^-selective response. Increased firing responses to rewarding cues: (A) Tone1, (B) Light1, and (F) Tone2+Light2. No increased firing occurred during presentation of unrewarding cues: (D) Tone2, (E) Light2, and (C) Tone1+Light1. Each filled circle below the raster line indicates one lick. Lower histogram shows summed licks. Onset of CS at time 0; negative values represent the pre-trial control. Each histogram bin, 80 ms. Suc, 0.3 M sucrose solution. Right vertical line calibrated firing rates by 10 spikes/s. This is the same neuron shown in Fig. 2A and B.

### Population coding of cue-reward associations

The response magnitudes of the 53 CS^+^-selective neurons were analyzed with multidimensional scaling (MDS; Fig. 4). The resulting r^2^ (0.975) and stress values (0.141) in the MDS analysis indicated that the responses to the CSs were well represented in 2D space. The MDS data indicated that there are two groups of CSs separated by a dotted line in Fig. 4: CSs associated with reward (Tone1, Light1, and Tone2+Light2) and CSs associated with nonreward (Tone2, Light2, and Tone1+Light1). The discriminant analysis indicated a significant separation between these two groups of the CSs (Wilks’ lambda = 0.031, p < 0.0001). Thus, this suggests that population activity of CS^+^-selective neurons codes reward values of the CSs. Furthermore, the MDS data indicated that there are six groups of CSs: Tone1, Light1, Tone2+Light2, Tone2, Light2, and Tone1+Light1. The multiple discriminant analysis indicated a significant separation among the six groups of CSs (Wilks’ lambda = 0.081, p < 0.0001). These data suggest that population activity of CS^+^-selective neurons separately represent the configural (Tone1+Light1 and Tone2+Light2) and elementary stimuli (Tone1, Light1, Tone2, and Light2); the population activity patterns distinguished the configural CSs from their constituents (elementary CSs), suggesting an RSC involvement in stimulus-stimulus association.

**Figure 4.**
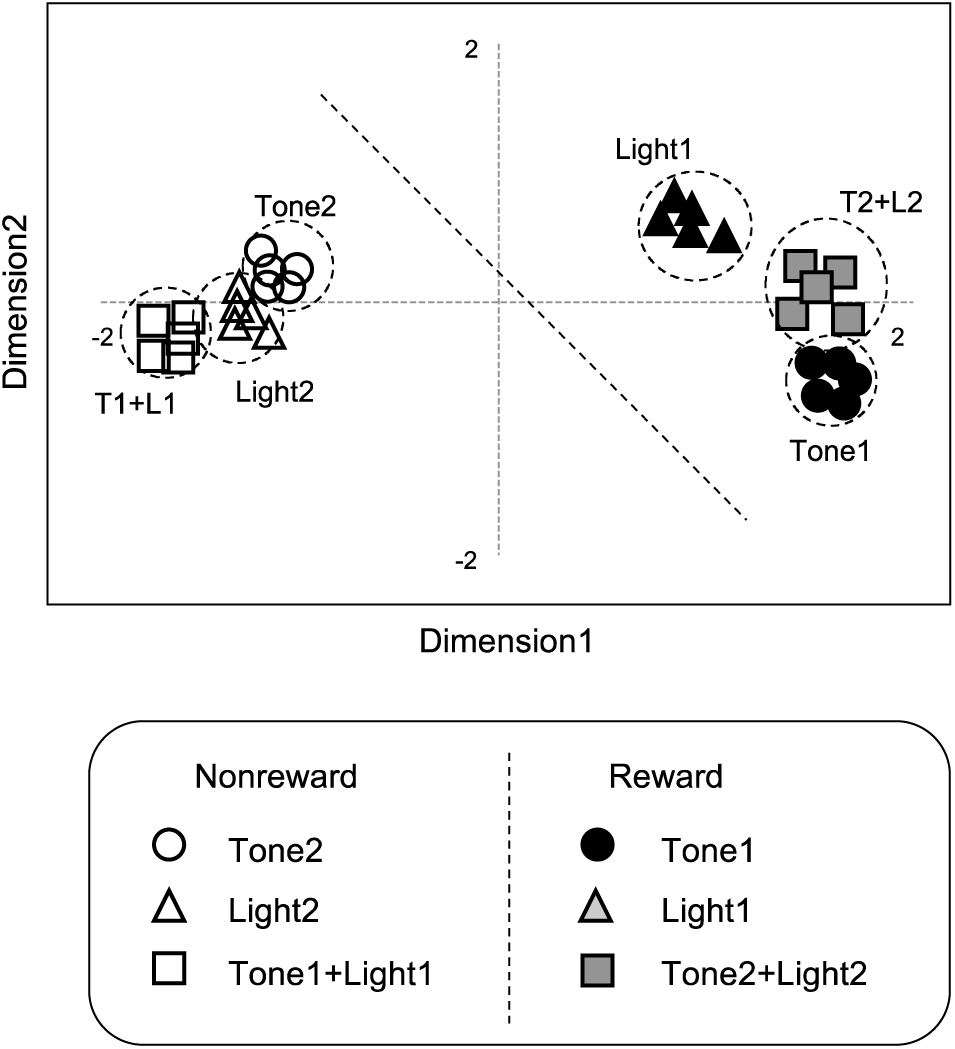
MDS analysis results showing distributions of the six CSs in two-dimensional space. Two clusters containing rewarding and unrewarding CSs as well as six clusters corresponding to six CSs were recognized.

### Theta rhythmicity of CS^+^-selective neurons during CS presentation

Theta rhythmicity in spike trains of CS^+^-selective neurons was analyzed by wavelet transform of spike trains. Figure 5A shows Morlet wavelet function used for wavelet transformation. Figure 5B shows an example of a CS^+^-selective neuron showing theta oscillation, where each spectral graph indicates mean spectra of each five trials of each CS. Spectral density in theta band increased during presentation of the CSs associated with reward (Fig. 5Ba, b, and f). Then, mean maximum theta band spectral power density during CS presentation across the 53 CS^+^-selective neurons were compared among the six CSs by one-way ANOVA. Figure 5C shows such comparisons of mean maximum spectral density in the theta band among the six CSs. The responses to the six cues had significantly different mean maximum spectral densities [Figure 5C; F(5, 312) = 13.114, p = 0.0001]. These were significantly greater for the rewarding CSs than the unrewarding ones (Bonferroni post hoc tests, all p’s < 0.01).

**Figure 5.**
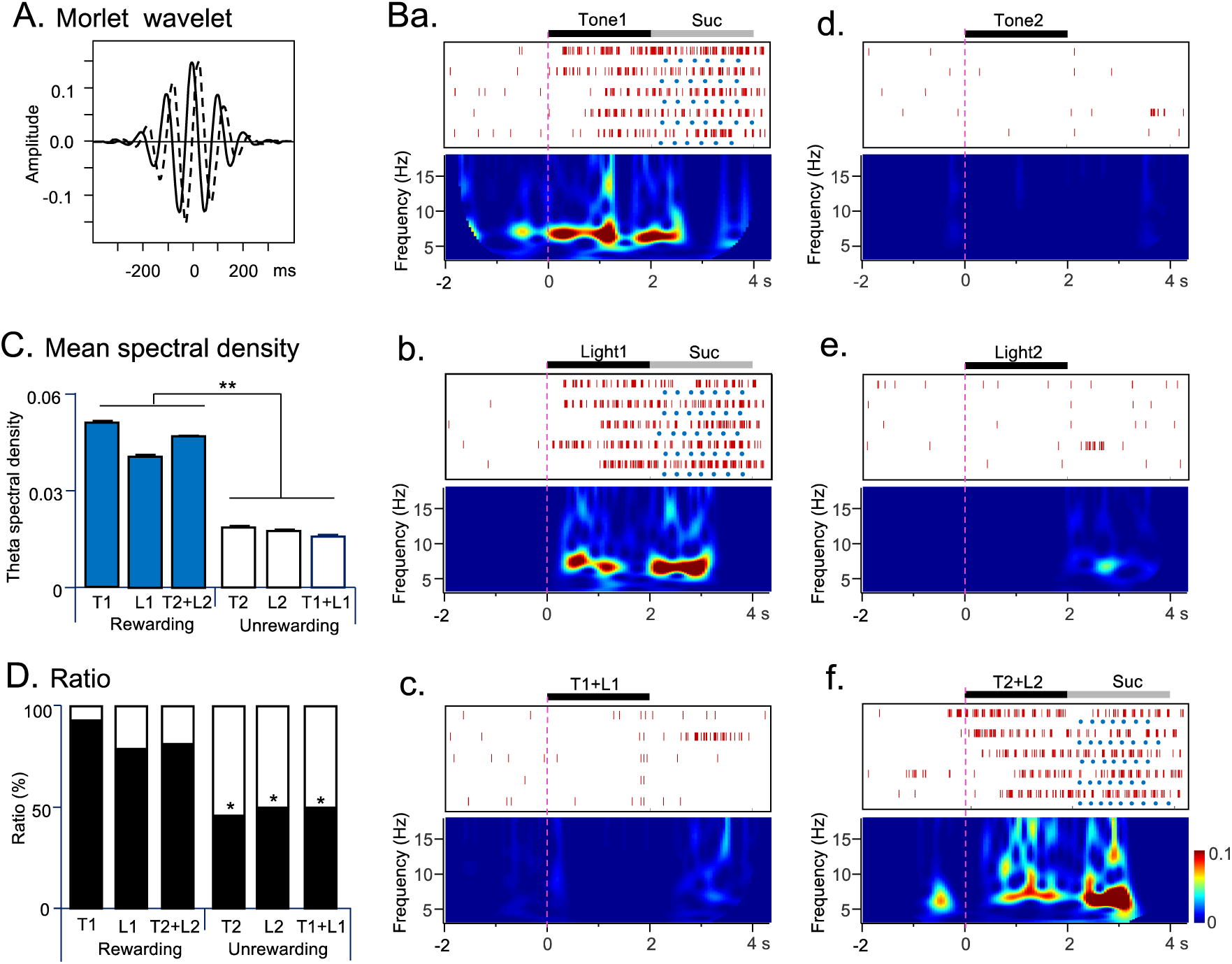
Rhythmic activity of CS^+^-selective neurons. A, Examples of Morlet wavelet functions (10 Hz) used for wavelet transformation. Solid and dotted lines indicate real and imaginary components, respectively. B, Example of wavelet spectra of a CS^+^-selective neuron. Increased theta oscillations in response to rewarding cues: (a) Tone1, (b) Light1, and (f) Tone2+Light2. No increased theta oscillations occurred during presentation of unrewarding cues: (d) Tone2, (e) Light2, and (c) Tone1+Light1. Onset of CS at time 0. C, Mean maximum theta band spectral density during presentations of the six CSs (n=53 neurons). **, p < 0.01 (Bonferroni test). D, Comparison of ratios of CS^+^-selective neurons with significant increases in maximum theta spectral density during presentation of each CS. Ratios of CS^+^-selective neurons with significant increases of theta oscillation were significantly greater in the rewarding CSs than the unrewarding CSs. Filled and open areas indicate ratios of CS^+^-selective neurons with significant and insignificant increases of theta oscillations in 53 CS^+^-selective neurons, respectively. *, significant difference from ratios in the rewarding CSs (McNemar post hoc test, p < 0.05).

We further analyzed significant changes in theta oscillations of neuronal firing during CS presentation compared with 2-s pre-CS period: significant increases in theta oscillation during each CS presentation were determined by comparison of maximum theta band (4-12 Hz) spectral power density during presentation of each CS with that during corresponding 2-s pre-CS period (Wilcoxon sum rank test, p < 0.05) in each CS^+^-selective neuron. Figure 5D shows ratios of CS^+^-selective neurons with significant increases in maximum theta spectral density during presentation of each CS in 53 CS^+^-selective neurons. There were significant differences in the ratios among the six CSs (Figure 5D; Cochran’s Q test, p < 0.0001). The ratios of CS^+^-selective neurons with significant changes in theta spectral density were significantly greater for the rewarding CSs than the unrewarding CSs (McNemar post hoc test, p < 0.05).

### Activity of CS^+^-selective neurons in error trials

The analyses in the above sections indicated that both firing rates and theta oscillation increased during presentation of the CSs associated with reward. During recordings of 18 CS^+^-selective neurons, in a few unrewarded trials, the rats erroneously licked the tube (error trials, eTrials). We hypothesized that if activity increases or increases in theta oscillations are involved in retrieval of memory of CS-reward association, incidental activity increases or increases in theta oscillation of CS^+^-selective neurons might occur during presentation of CSs associated with nonreward, which leads to licking. To test this idea, mean firing rates during presentation of the CS in eTrials were compared with those during presentation of the same CS in correct trials (cTrials) in 18 CS^+^-selective neurons (Fig. 6A). The results indicated that mean firing rates during presentation of the unrewarding CSs were significantly greater in the error trials than the correct (no lick) trials (paired t-test, p = 0.0077). Furthermore, theta oscillations of 18 CS^+^-selective neurons in a wavelet spectral analysis were similarly compared between the eTrials and cTrials (Fig. 6B). The results indicated that mean maximum spectral density during presentation of the unrewarding CSs were significantly greater in the error trials than the correct trials (paired t-test, p = 0.00021). These results suggest that activity increases with theta oscillation of the CS^+^-selective neurons during presentation of CSs are associated with reward anticipation due to incorrect retrieval of memory for rewarding CSs.

**Figure 6.**
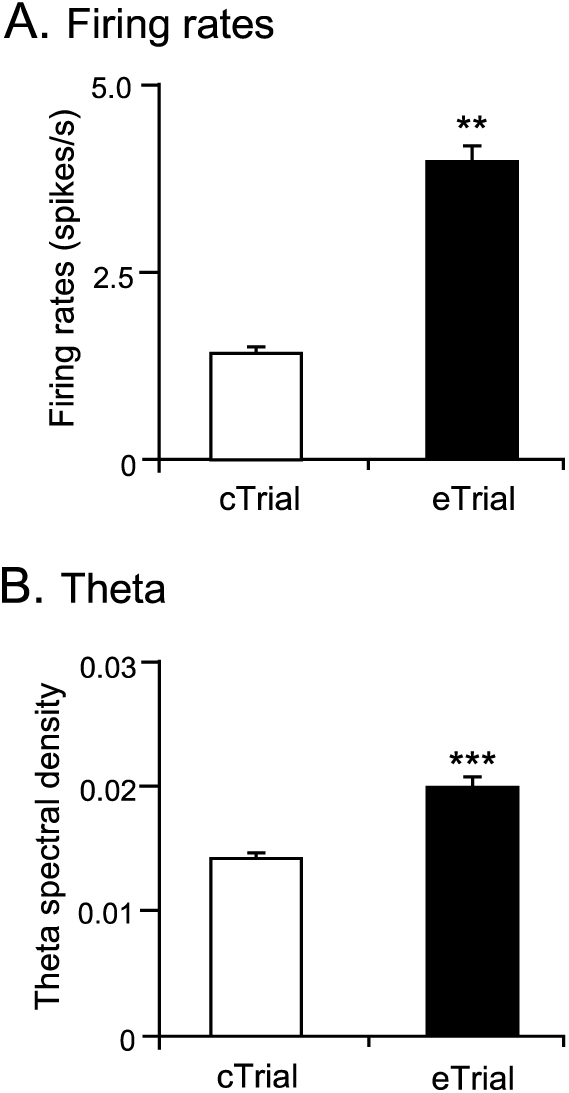
Comparison of mean firing rates (A) and mean maximum theta spectral density (B) between the correct (cTrials) and error (eTrials) trials during presentation of unrewarding CSs in CS^+^-selective neurons. **, ***, p < 0.01, 0.001, respectively.

### Phase locking of spikes to theta

Phase-locking of neuronal spikes of 53 CS^+^-selective neurons relative to theta rhythms (4-12 Hz) in the LFPs recorded from the same electrode, where neuronal activity was recorded, was separately analyzed during presentations of the rewarding and unrewarding CSs. An example of such phase-locking to theta cycles is shown in Fig. 7. This neuron was phase-locked to theta cycles during presentation of rewarding and unrewarding CSs. The mean phase-locking angle was 260°, near the trough of the theta cycle for rewarding and unrewarding cues (arrows in Fig. 7Aa,b and Ba,b).

**Figure 7.**
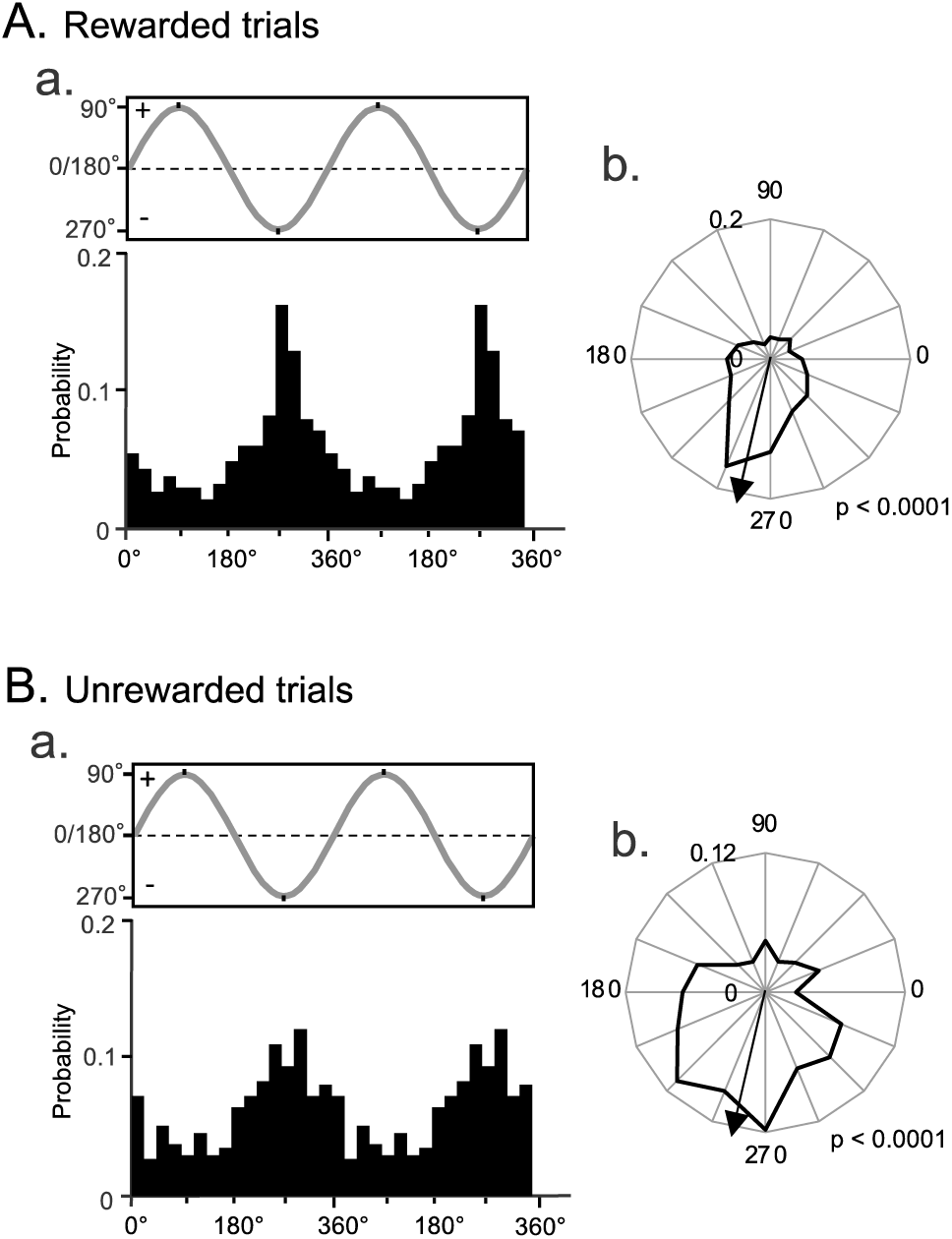
Spike phase locking to LFP theta in the rewarded (A) and unrewarded (B) trials. (a) Firing probability of this neuron as a function of LFP theta phase. (b) Polar plots of preferred phases of spike train (for A and B, Rayleigh’s test, p < 0.0001). Values at the top left of the polar plots indicate ratios of spikes within 22.5° width to the total number of spikes. Arrows indicate phase locking angles.

Figure 8A shows mean PLV of CS^+^-selective neurons with such significant phase locking to LFP theta cycles. The mean PLV was significantly greater in the rewarding CSs than the unrewarding CSs (n=14; paired t-test, p=0.0033) (Fig. 8A). Figure 8B shows the distribution of mean phase-locking angles across 53 CS^+^-selective neurons. In Fig. 8Ba, the data during presentation of the rewarding and unrewarding CSs were separately analyzed. Since previous studies suggest that phase locking to the positive and negative phases of the theta LFP cycles are associated with encoding and retrieval, respectively (Hasselmo, 2002; Cutsuridis et al., 2010; Douchamp et al., 2013), CS^+^-selective neurons were divided into two subgroups based on the mean phase-locking angles; the CS^+^-selective neurons with mean phase-locking angles in the trough (negative) phase (180-360°) and those with mean phase-locking angles in the peak (positive) phase (0-180°). Ratios of the mean phase-locking angles in the negative phase in 53 CS^+^-selective neurons were significantly greater than those in the positive phase during presentation of the rewarding CSs (χ^2^-test, p=0.0038). In response to the unrewarding CSs similar results were obtained: ratios of the mean phase-locking angles in the negative phase were significantly greater than those in the positive phase (χ^2^-test, p < 0.0001). When data during presentation of both rewarding and unrewarding CSs were combined (Fig. 8Bb), comparable results were obtained: ratios of the mean phase-locking angles distributed in the negative phase were significantly greater than that in the positive phase during presentation of the CSs (χ^2^-test, p < 0.0001). These results indicate that the CS^+^-selective neurons preferentially fired near the trough of the LFP theta cycles.

**Figure 8.**
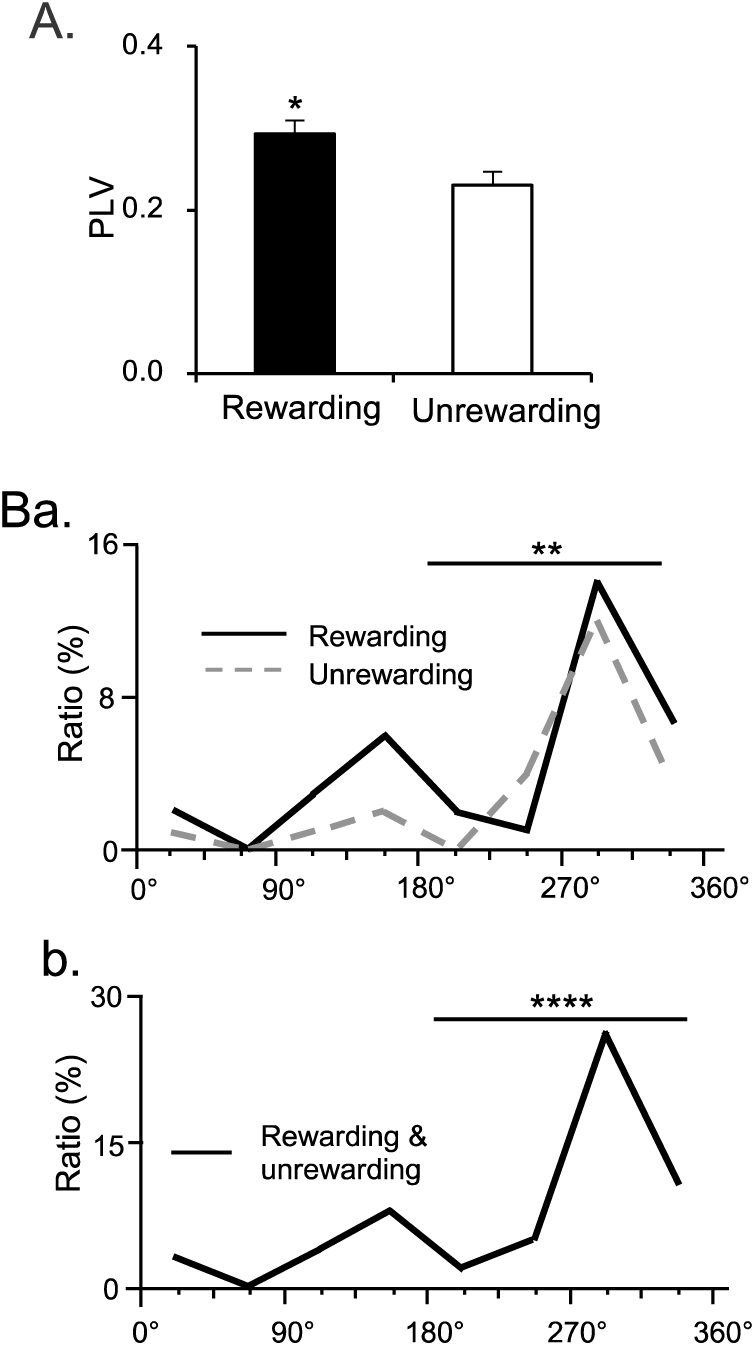
Spike-LFP coupling (A) and distribution of mean phase-locking angles (B) of the CS^+^-selective neurons. A, Comparison of PLV during presentation of the rewarding and unrewarding CSs. B, Angle distribution was analyzed using separate data sets during presentation of rewarding and unrewarding CSs (a) and combined data during presentation of the all CSs (b). Ratios of CS^+^-selective neurons with mean phase-locking angles in the negative phase between 180-360° were significantly greater than those in the positive phase between 0-180°. *, **, ****, p < 0.05,

### Locations of the RSC neurons

The distributions of CS^+^-selective neurons, CS-nondifferential neurons, other differential neurons, and nonresponsive neurons are shown in Figure 9A-C, respectively. The various types of RSC neurons were located in both the superficial (layers 2-3) and deep (layers 4-5) layers of the RSC.

**Figure 9.**
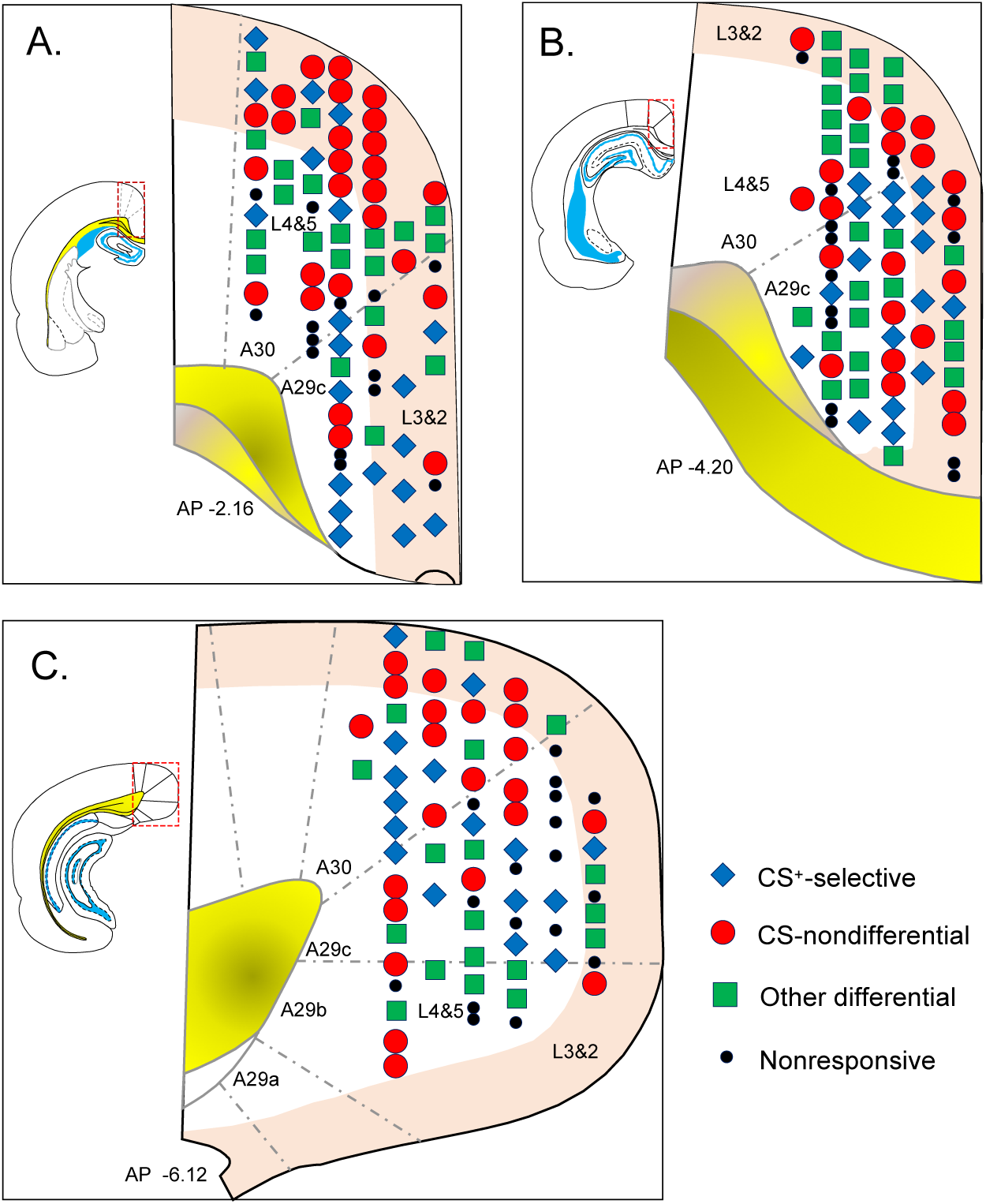
Locations of the recording sites of RSC neurons. A-C, neurons are plotted on coronal sections of the RSC. The rectangular areas surrounded by the dotted lines in the left brain sections are enlarged on the right. Numbers below brain sections indicate the distance (mm) posterior from the bregma. ‘L2&3’ with colored background: layers 2 and 3; ‘L4&5’ without the background: layers 4 and 5.

## Discussion

### Sensory coding in the RSC

Here, a group of 53 out of 234 RSC neurons responded selectively to multimodal CSs associated with reward. In well-trained animals, the RSC neurons responded stronger to Tone1 and Light1 associated with reward than Tone1+Light1 associated with nonreward. Thus, the neurons discriminated the compound negative pattering stimulus (i.e., Tone1+Light1) from the elementary stimuli (i.e., Tone1 and Light1). Similarly, these neurons responded more strongly to Tone2+Light2 associated with reward than Tone2 and Light2 associated with nonreward. These results indicate that the neurons discriminated the compound positive patterning stimulus (i.e., Tone2+Light2) from its elementary stimuli (i.e., Tone2 and Light2). In addition, the MDS analyses indicated that the neural representations of elementary stimuli were distinct from their compound stimuli (i.e., Tone1 and Light1 vs. Tone1+Light1; Tone2 and Light2 vs. Tone2+Light2). These results support previous behavioral studies showing that the RSC is involved in stimulus-stimulus association to discriminate configural and elemental stimuli. Previous behavioral studies with RSC lesions and silencing of RSC neurons also reported an RSC involvement in stimulus-stimulus association (Gibbs and Johnson et al., 2007; Keene and Bucci, 2008b; Robinson et al., 2011; Robinson et al., 2014; Bucci and Robinson, 2014; Fournier et al., 2019a). Anatomical connections of the RSC also support this idea: the RSC is considered to be a hub that receives multimodal (including auditory and visual) projections to integrate diverse information (see Introduction). Taken together, the present results provide a neurophysiological basis of an RSC role in stimulus-stimulus association. However, note that we recorded the RSC neurons from well-trained rats, so they might actually be involved in storing and expression of remote memory of stimulus-stimulus associations (discussed in the next section).

Here, the RSC neurons responded stronger to the CSs associated with reward than the CSs associated with nonreward. These findings are consistent with previous studies reporting that multi-unit activity in the rabbit area 29 responded stronger to CSs associated with rewards (Gabriel et al., 1987; Freeman et al., 1996; Smith et al., 2004). Furthermore, Vedder et al. (2017) reported that during learning the ratios of RSC neurons responsive to cues important for task performance increased. These findings suggest that the RSC neurons may code “significant” stimuli important for animals to perform tasks (Smith et al., 2018). Consistent with this idea, the human RSC has been reported to be consistently activated in response to emotionally salient stimuli (Maddock, 1999). A previous study suggests that neuronal responses to CSs were modulated in the primary visual cortex according to top-down signals from the RSC (Makino and Komiyama, 2015). The MDS analyses showed separate representations for the CSs associated with reward and those associated with nonreward, with elementary cues and configural cues grouped together. The present results suggest that this significance information for each CS represented in the RSC function as top down signals.

### Role of the RSC in retrieval of remote memory

Here RSC neurons responded to nonspatial CSs in well-trained rats. These neuronal activities might reflect retrieval of remote memory stored in the RSC. Consistent with this, human imaging studies have reported that the RSC was active during retrieval of past memories (Shannon and Buckner, 2004; Daselaar et al., 2006; Svoboda et al., 2006), while RSC damage induced not only anterograde amnesia, but also retrograde amnesia in humans (Valenstein et al., 1987). Furthermore, RSC lesions or pharmacological inactivation (e.g., chemogenetic inactivation, injection of muscimol or NMDA blocker) before retention tests induced deficits in retrieval of long-term memory in rodents (Corcoran et al., 2011; Haijima and Ichitani, 2012; Katche et al., 2013; Todd et al., 2016; Jiang et al., 2018; Fournier et al., 2019b). In addition, stimulus-stimulus association in the RSC might underlie coding of contexts (or places), which consist of multiple stimuli or objects (Smith et al., 2018). Indeed, optogenetic activation of RSC neurons induced same freezing behaviors as those observed by presentation of a contextual stimulus associated with shock (Cowansage et al., 2014). The present results, along with the above previous studies, are consistent with role of RSC in storing and expression of remote memory.

Studies in humans reported that theta oscillations were associated with retrieval of long-term memory (Nyhus and Curran, 2010; Burke et al., 2014; Nyhus et al., 2019): theta power was increased in response to items that subjects previously studied compared to those that the subjects previously had not encountered. Furthermore, a recent study reported that theta oscillations occurred in the RSC 500-200 ms before retrieval of items that subjects previously studied (Kerrén et al., 2018). Therefore, we further analyzed oscillatory activity of RSC neurons. The CS^+^-selective neurons showed stronger theta oscillations during presentation of the rewarding CSs compared to the unrewarding CSs. Furthermore, both neuronal firing rates and theta oscillation of the CS^+^-selective neurons increased in those error trials when the rats licked the tube in response to the unrewarding CSs. These results suggest that increases in neuronal activity and theta oscillations in the RSC are essential for expression and retrieval of remote memory of the rewarding CSs. The present results also indicated that neuronal activity of the CS^+^-selective neurons was phase-locked to theta rhythm of LFPs in the RSC. Furthermore, more CS^+^-selective neurons fired in the trough of LFP theta rhythm in the present study. LFP theta rhythm in the RSC is in-phase with that in the hippocampal CA1 (Young and McNaughton, 2009; Koike et al., 2017). It has been proposed that CA1 neurons fire in the trough of the theta rhythm when animals recall previous memory (Hasselmo, 2002; Cutsuridis et al., 2010; Douchamp et al., 2013). These findings suggest that activity of RSC neurons might be synchronous with CA1 neuronal activity during retrieval of remote memory. Human studies reported that EEG theta oscillations occurred before retrieval of remote memory or learned associative memory (Burke et al., 2014; Kerrén et al., 2018), suggesting that they might reflect memory search and information transfer between the hippocampal formation and the cortical areas such as the RSC (Fell and Axmacher, 2011; Burke et al., 2014). The hippocampal formation has been implicated in a retrieval process in which, when partial information is transferred to the hippocampal formation, entire memory is represented through hippocampal projections (“index”) to cortical area related to entire memory to which the partial information belongs (Teyler and Rudy, 2007). The information transfer from the RSC to the hippocampal formation may promote this process in retrieval of remote memory (Bucci and Robinson, 2014; Miller et al., 2014). The present study provides neurophysiological evidence in the RSC underling these retrieval processes of remote associative memory.

## Materials and Methods

### Subjects

Five male albino Wistar rats, weighing 250–300 g (10–18 weeks old; SLC, Hamamatsu, Japan), were used. The rats were individually housed in cage (temperature 23 ± 1°C; 12-h light–dark cycle; and free access to chow and water). Housing conditions, care and treatment of the animals during all stages of the experiments conformed to the National Institutes of Health on the Care of Humans and Laboratory Animals and the Guidelines for the Care and Use of Laboratory Animals at the University of Toyama. The study was approved by the Ethical Committee of Animal Experiments at the University of Toyama (Permit No.: A2014MED-37 and A2017MED-16).

### Surgery

As in our previous studies (Uwano et al., 1995; Nishijo et al., 1998; Munkhzaya et al., 2020; Chinzorig et al., 2020), we used our head restraint system associated with a stereotaxic instrument (Nishijo and Norgren, 1990, 1991, 1997) (see Figure 1A). For this, briefly, the rats were anesthetized with an anesthetic mixture of midazolam (2 mg/kg, i.p.), medetomidine (0.38 mg/kg, i.p.), and butorphanol (2.5 mg/kg, i.p.), and five small, sterile, stainless screws were implanted into the skull to serve as anchors for dental acrylic. Then, the cranioplastic acrylic was built up on the skull and molded around the conical ends of two sets of stainless-steel bars. Once the cement had hardened, these bars were removed, leaving a negative impression on each side of the acrylic block. These artificial earbars were used to painlessly hold their heads in the stereotaxic instrument (Figure 1A). A short length of 27-gauge stainless-steel tubing was embedded into the cranioplastic acrylic near the bregma on the skull to serve as a reference pin during chronic recording. After surgery, an antibiotic was administered.

After recovery from surgery (10–14 days) and training (2 weeks), a hole (diameter, 2.8– 3.0 mm) was drilled through the cranioplastic and the underlying skull (A, −1.72 to −6.6 mm from bregma; L, 0.3 mm left and right) for semi-chronic recording under anesthesia with an anesthetic mixture of midazolam, medetomidine, and butorphanol (see above). The exposed dura was removed, and one or two drops of chloramphenicol (Sankyo Co., Ltd., Tokyo, Japan) solution (0.1 g/ml) were administered into the hole. The hole was covered with a sterile Teflon sheet and sealed with epoxy glue. After the animal recovered (5–7 days), it was placed back on the water-deprivation regimen.

### Task paradigms and training

Task paradigms and training were essentially similar to our previous studies (Oyoshi et al., 1996; Takenouchi et al., 1999; Toyomitsu et al., 2002; Matsuyama et al., 2011; Munkhzaya et al., 2020). Briefly, rats were maintained under a 12-h water-deprivation regimen. The ear bars of a stereotaxic instrument were fixed to the molded dental acrylic on the headstage acrylic and the restrained rat was placed in front of the behavioral apparatus (Figure 1A). The rats were required to discriminate among six CSs associated with reward (0.3 M sucrose solution) or nonreward (Table 1) in the cue-reward association task. A speaker located 50 cm in front of the rat delivered the auditory stimuli, and two white lights, 3 cm in front of each eye, served as visual stimuli. Licking was detected by a touch sensor on the tube. Reward availability from the tube was signaled by either a 2860 Hz continuous tone (Tone 1, an elemental stimulus), the right light cue (Light 1, an elemental stimulus), or the simultaneous presentation of a 530 Hz tone (Tone 2) and the left light cue (Light2) (Tone2+Light2; a configural stimulus). The tube was extended at the end of the stimulus, and the rat could obtain reward if it licked the tube during 2 s period after these cues (rewarded trials: Figure 1Ba). Tone2, Light2, or simultaneous presentation of Tone1 and Light1 (Tone1+Light1; configural stimulus) signaled nonreward (unrewarded trials: Figure 1Bb).

The rats were initially trained to lick a tube, which was automatically extended close to the mouth for 2 s to deliver reward. Training in the rewarded or unrewarded trials was then conducted in one block of 15 or 20 trials for each cue. In the block trials with nonreward, the rats learned not to lick the dry tube. After the rats learned to respond to each cue in the block trials, they were trained to discriminate cues presented in pseudo-random sequence until correct performance exceeded 90%. Throughout the training and recording periods, rats were permitted to ingest 15–30 ml of sucrose solution while under restraint. If the rat failed to consume 30 ml of the solution while restrained, it was given the remainder when it was returned to its home cage. After the rats reached criterion performance, RSC neurons were recorded while they performed the cue-reward association task.

### Electrophysiological recordings

Each rat was usually tested every other day. After the head was placed in the stereotaxic apparatus, a glass-insulated tungsten microelectrode (impedance 1.0–1.5 MΩ at 1 kHz) was stereotaxically inserted stepwise with a pulse motor-driven manipulator (SM-20, Narishige, Tokyo, Japan) into the RSC region. During recording, each CS was pseudo-randomly presented with an inter-trial interval of 20-30 s. Both single-unit spikes and LFPs were recorded simultaneously. Spike activity was filtered from 0.1-8 kHz. LFP signals were filtered from 3.3 to 88 Hz with first order high- and low-pass filter circuits. Signals were referenced to a skull screw over the cerebellum. The spike and LFP signals, and the lick contacts on the tube were digitized and stored with the cue trigger signals in a personal computer through a Multichannel Acquisition Processor (MAP, Plexon Inc., Dallas, TX, USA) system. The sampling rates of the spike and LFP signals were 40 kHz and 1 kHz, respectively.

Spikes were sorted into single units with the Offline sorter program (Plexon Inc.) for cluster analysis. For each isolated cluster, an autocorrelogram was constructed and only units with refractory periods > 2.0 ms were analyzed further. Waveforms of the isolated units were superimposed to check the consistency of the waveforms, and then they were transferred to the NeuroExplorer program (Nex Technologies, Madison, USA) for further analysis.

### Data analyses

#### Analysis of RSC neuronal activity in the task

Each CS was usually tested in 5-8 trials for each neuron. We analyzed single neuronal activity during the 2 s CS period and 2 s tube protrusion period in the last five correct trials. The baseline firing rate was defined as the mean firing rate during the 1-s pre-CS period. Significant excitatory or inhibitory responses to the CSs were defined by Wilcoxon sum rank test (p < 0.05) of neuronal activity of mean firing rate between the 1-s pre-CS and the 2 s CS periods. RSC neurons with significant responses at least to one CS were defined as CS-responsive neurons. The response magnitudes to the six CSs were further analyzed by one-way ANOVA (p<0.05) with a Bonferroni post-hoc test (p<0.05). The response magnitude to CSs was defined as ([mean firing rate during CS period for 2 s] minus [mean firing rate during the pre-CS period for 1 s]). The RSC neurons with no significant main effect were defined as CS-nondifferent neurons, while the RSC neurons with a significant main effect were defined as CS-differential neurons (Table 2). The CS-differential neurons were further divided into two subgroups based on post hoc tests. If response magnitudes to all three CSs associated with rewards were greater than those associated with non-reward, they were considered CS^+^-selective neurons. The remaining differential neurons were defined as other CS-differential neurons. Significant excitatory or inhibitory responses during the tube protrusion period after each CS were similarly defined with a Wilcoxon sum rank test (p < 0.05) comparing mean firing rate between the 1-s pre-CS and the tube protrusion periods.

#### Population coding analyses

The response magnitudes of CS^+^-selective neurons were further analyzed by multidimensional scaling (MDS) analysis. Each of these neurons was repeatedly (i.e., five times) tested with six CSs. This yielded 30 stimulus arrays. Then, the data matrices of the response magnitudes in the 53 × 30 arrays derived from the 53 CS^+^-selective neurons were generated. The Euclidean distances of the dissimilarities between all possible pairs of stimuli were calculated. The MDS program (PROXSCAL procedure, SPSS Statistical Package, version 16; IBM Corporation, NY, USA) positioned the visualization in a two-dimensional (2D) space with the distances between the stimuli representing the original relationships (i.e., the Euclidean distances in the present study) (Shepard, 1962). The clusters of the CSs were evaluated using the multiple discriminant analysis.

#### Theta modulation analyses

Theta rhythmic firing (4-12 Hz) of CS^+^-selective neurons was analyzed by taking the wavelet transform of their spike trains (Lee, 2003) with the FieldTrip MATLAB toolbox (Oostenveld et al., 2011). Mean maximum theta band spectral power density during CS presentation across the 53 CS^+^-selective neurons were compared among the six CSs by one-way ANOVA with post hoc test (Bonferroni test, p < 0.05). Furthermore, significant changes in theta oscillation during CS presentation were determined by comparison of maximum theta band (4-12 Hz) spectral power density during presentation of each CS with that during corresponding 2-s pre-CS period (Wilcoxon sum rank test, p < 0.05) in each CS^+^-selective neuron.

#### Phase-locking analyses

LFPs were initially filtered to the theta band (4-12 Hz) (NeuroExplorer). Phase-locking of neuronal spikes of 53 CS^+^-selective neurons relative to theta rhythms (4-12 Hz) in the LFPs recorded from the same electrode, where neuronal activity was recorded, was separately analyzed during presentations of the rewarding and unrewarding CSs. Phase-locking of spikes to LFP theta cycles was analyzed with the Freely Moving Animal MATLAB Toolbox (http://fmatoolbox.sourceforge.net/). Theta phase (with 270° as the trough), to which spikes were phase-locked, was tallied with 22.5° bin widths. Significance of phase-locking was tested by Rayleigh test (p < 0.05). Degree of spike-LFP coupling with significant phase locking was evaluated by computing phase locking value (PLV), as follows:

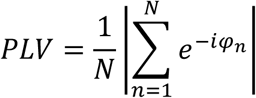

where *N* represents total number of spikes and *φ*_*n*_ represents the phase of LFP at which *n*-th spike occurs. Since PLV is dependent on total number of spikes (Zarei et al., 2018), only neurons with total numbers of spikes greater than 50 during presentation of rewarding and unrewarding CSs were analyzed.

A mean phase angle of phase locking was estimated using the above software in each CS^+^-selective neuron with significant phase locking, and further analyzed. The CS^+^-selective neurons were divided into two subgroups based on the mean angles; the CS^+^-selective neurons with mean angles in the trough phase (180-360°) and those with mean angles in the peak phase (0-180°). Significant bias of mean angles was tested by Chi-square test (p < 0.05) of the ratios of these two types of CS^+^-selective neurons in 53 CS^+^-selective neurons.

### Histological analysis of recording sites

After the last recording, the rats were deeply anesthetized with sodium pentobarbital (100 mg/kg, i.p.) and several small electrolytic lesions (20 μA for 20 s) were made through electrodes re-inserted at the previously recorded sites. The rats were then removed from the apparatus and perfused transcardially with 0.9% saline and 10% buffered formalin. The coronal brain sections (50 μm) were stained with Cresyl violet. Recording positions of neurons were stereotaxically located on the actual tissue sections and plotted on the corresponding sections of the Paxinos and Watson atlas (2017).

## Acknowledgements

We thank Dr. S. Wiener (CNRS) for valuable comments on the manuscript. The study was supported partly by Takeda Science Foundation and research grants from University of Toyama.

## Source data files

Figure 2-source data 1. Source data in Figure 2B.

Figure 2-source data 2. Source data in Figure 2C.

Figure 4–source data 1. Source data in Figure 4.

Figure 5–source data 1. Source data in Figure 5C.

Figure 5–source data 2. Source data in Figure 5D.

Figure 6–source data 1. Source data in Figure 6A.

Figure 6–source data 2. Source data in Figure 6B.

Figure 7–source data 1. Source data in Figure 7Ab.

Figure 7–source data 2. Source data in Figure 7Bb.

Figure 8–source data 1. Source data in Figure 8A.

Figure 8–source data 2. Source data in Figure 8Ba.

Figure 8–source data 3. Source data in Figure 8Bb.

